# An EOMES induced epigenetic deflection initiates lineage commitment at mammalian gastrulation

**DOI:** 10.1101/2023.03.15.532746

**Authors:** Chiara M. Schröder, Lea Zissel, Sophie-Luise Mersiowsky, Mehmet Tekman, Simone Probst, Katrin M. Schüle, Sebastian Preissl, Oliver Schilling, H. Th. Marc Timmers, Sebastian J. Arnold

## Abstract

Summary paragraph

Different cell types are determined by cell lineage-specific transcriptional programmes and by epigenetic regulation of chromatin^1, 2^. Yet, the functional relationships between dynamically expressed transcription factors (TFs) and chromatin changes guiding lineage specification often remain elusive^3^. First mammalian embryonic lineages segregate when pluripotent cells become committed to either Mesoderm and Endoderm (ME) or Neuroectoderm (NE). NE forms by default in the absence of signalling-induced ME specification^4, 5^, resulting from global asymmetries in chromatin state favouring NE gene programme activation as recently demonstrated^6–8^. In this study, we unravel the initiation of ME lineage specification by the genome-wide, *de novo* formation of chromatin accessibility at ME enhancers that epigenetically deflects pluripotent cells from default NE differentiation. The Tbx TF *Eomes*, previously considered a transcriptional regulator, acts as global chromatin organizer that establishes ME lineage competence. EOMES recruits the canonical ATP-dependent chromatin remodelling complex SWI/SNF to broadly generate the chromatin-accessible ME enhancer landscape. This lineage competence is generated independently of ME gene transcription that fully depends on ME-inducing signalling pathways including Wnts and TGFβ/NODAL^9^. This study thus resolves the successive steps of ME lineage differentiation by globally establishing chromatin accessibility for lineage competence, followed by signal-encoded transcriptional regulation of different ME lineage-defining gene programmes.

## Main Text

During mammalian embryogenesis, the first embryonic lineage segregation of pluripotent cells generates ME and NE progenitors with gross differences in lineage-specific transcriptional outputs^10, 11^. Classic studies performed more than two decades ago demonstrated that pluripotent cells differentiate by default towards NE cell types in the absence of secreted, instructive signals for ME lineage specification, including BMPs, TGFβ and Wnts^5, 9, 12, 13^. More recent molecular analyses revealed that ME instructive signals are relayed by activities of the two Tbx TFs *Eomes* and *Brachyury* (a.k.a. *T*)^8^. The combined deletion of both Tbx factors in pluripotent mouse embryonic stem cells (mESCs) abolishes ME lineage potential, and consequently *Eomes* and *Brachyury* double-deficient (dKO) mESCs follow the default path to NE lineages in the presence of ME-inducing signals^8^. The molecular basis for this NE-directed lineage preference lies in preformed asymmetries in the chromatin landscape between NE and ME programme-associated genes in pluripotent cells. Cis-regulatory elements (CREs) of early NE-associated genes are generally chromatin accessible and hypomethylated, while the CREs of ME genes are inaccessible and hypermethylated in pluripotent cells^6–8^.

To date, neither the molecular regulation, nor the precise temporal order of chromatin remodelling processes that deflect pluripotent cells from the preformed NE fate bias during ME lineage specification are known. Thus, it is unclear if the broad changes of the chromatin landscapes found in differentiated ME cells reflect an early prerequisite, or rather occur as a consequence of ME lineage differentiation^8^.

### Chromatin dynamics during ME specification

First, to reveal the temporal dynamics of chromatin changes during the initial steps of ME lineage specification, we used *in vitro* differentiation of mESCs as 3D embryoid bodies (EBs) and induced ME differentiation by treatment with the NODAL analogue Activin A (ActA) (Fig. 1a)^8^. This protocol reflects specification of the first epiblast-derived lineages anterior mesoderm (AM) and definitive endoderm (DE)^8^. We charted chromatin accessibility changes of wild type (WT) EBs by ATAC-seq (Assay for Transposase-Accessible Chromatin using sequencing) and profiled transcriptional changes by RNA-seq at 12 hrs intervals starting from day 2 (d2) after aggregation at EPI stem cell state^14^, and monitored early ME lineage specification until d3.5 (Fig. 1a). ATAC-seq resulted in a total of 37,846 accessible sites (FDR≤0.05) collected across the four time-points from d2 to d3.5 (Supplementary Table 1). Remarkably, of all ATAC-sites ∼72% (n=27,088 peak sites) show significant changes in peak intensities within this 36 hrs time-period (Fold>1 or Fold<-1, FDR≤0.05; Extended Data Fig. 1a, Supplementary Table 1), indicating the highly dynamic, global regulation of chromatin accessibility during early ME differentiation.

**Fig. 1.**
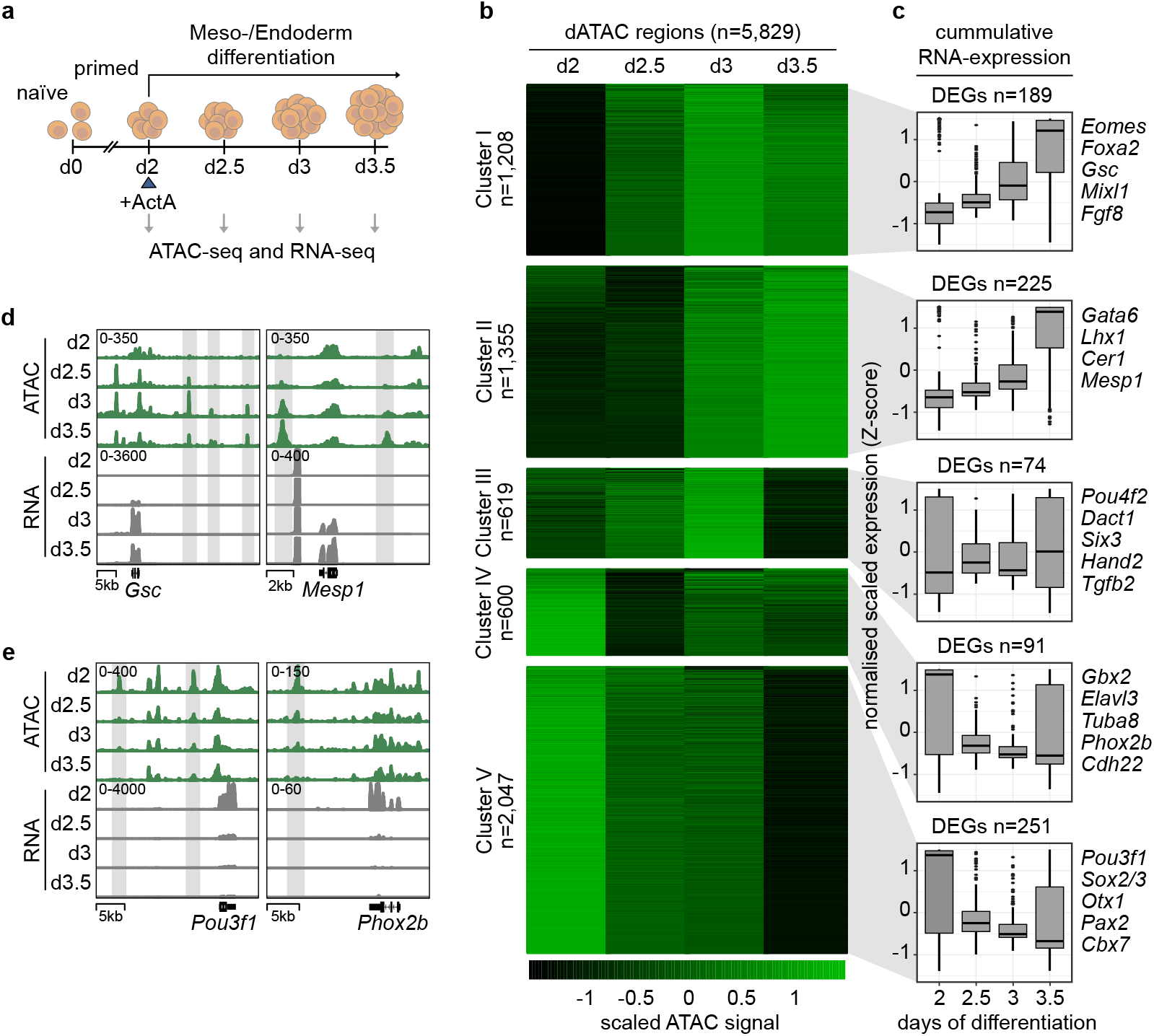
Global dynamics of chromatin accessibility during ME differentiation. **a**, Schematic of differentiation to Mesoderm and Endoderm (ME) by aggregation of naïve mouse embryonic stem cells (mESCs) as embryoid bodies (EBs) in basal, serum-free medium from day 0 (d0) to d2, and ME induction by 30 ng/ml Activin A (ActA). Samples were collected in 12 hours intervals for ATAC-seq and RNA-seq analysis from d2 (primed state) to d3.5. **b**, Heat map showing the dynamic changes in chromatin accessibility during ME specification. The 10,000 most dynamic ATAC (dATAC) sites (of all ATAC sites detected between d2-d3.5) were associated to genes, and sites that converge on the same gene were merged resulting in n=5,829 dATAC regions. Regions were clustered by K-means into five indicated clusters according to the dynamics of accessibility changes, e.g. early (I), late accessible (II), transient (III) accessible, and early (IV), late (V) inaccessible sites. **c**, Box-whisker plots showing cumulative expression of differentially expressed genes (DEGs) within each dATAC cluster. Numbers of DEGs within each dATAC cluster are indicated. The y-axis shows the averaged, normalised scaled expression (Z-score) of DEGs between d2-d3.5 of differentiation (indicated on x-axis). The 25th and 75th percentiles are represented in the boxes; the horizontal black line indicates the median. Whiskers extending no more than 1.5 times the inter-quartile range (IQR). Outliers are indicated by circles. **d**, Genome browser views of ATAC-seq (green) and RNA-seq (grey) tracks of definitive endoderm (*Gsc,* cluster I) and anterior mesoderm (*Mesp1,* cluster II) marker genes, showing the dynamic increase in accessibility, and **e**, of anterior epiblast/NE genes *Pou3f1* (cluster V), and *Phox2b* (cluster IV) showing decreased accessibility and loss of expression (grey box). Tracks show normalised counts to RPKM from 2 or 3 merged replicates.

Comparing chromatin accessibility changes between consecutive time points showed that between d2-d2.5 accessible sites are predominantly closing, followed by a rapid increase of accessibility between d2.5-d3 (Extended Data Fig. 1a). For the more detailed analysis of these highly dynamic chromatin accessibility changes we selected 10,000 most variable sites that we refer to as dynamic ATAC (dATAC) sites (Supplementary Table 2). dATAC sites were associated to nearest genes and multiple sites merged when they converged onto the same gene, resulting in n=5,829 gene-associated dATAC regions (Fig. 1b, Extended Data Fig. 1b). K-means clustering of dATAC regions resulted in five clusters that relate to the dynamic patterns of opening and closing sites during differentiation (Fig. 1b, Supplementary Table 3). Cluster I shows early (d2.5) and cluster II delayed (d3) gain of accessibility, cluster III regions are transiently accessible, and regions of clusters IV and V gradually loose accessibility (Fig. 1b, Supplementary Table 3).

To integrate chromatin accessibility dynamics with transcriptional changes, we clustered the n=2,297 differentially expressed genes (DEGs, adjusted P≤0.05, log_2_(FC)≥2) of RNA-seq data between d2-d3.5 into five clusters of genes with dynamic expression changes (Extended Data Fig. 1c, Supplementary Table 4). Clusters 1-3 contained prominent ME genes that follow a temporal expression pattern of transient (cluster 1), early upregulated (cluster 2) and delayed upregulated ME genes (cluster 3). Clusters 4 and 5 are characterised by the gradual decrease in expression and contain known markers of pluripotency and anterior epiblast/early NE fate (Extended Data Fig. 1c). We calculated cumulative, normalised expression values of the DEGs that associated to the five dATAC clusters (n=830, Fig. 1c) and found that these follow the pattern of chromatin accessibility dynamics of the CREs (Fig. 1c), such that accessibility at ME marker genes increases together with expression (DEGs at clusters I and II), and the expression levels of anterior epiblast/early NE and pluripotency genes are gradually decreased following the reduction of accessibility (DEGs at clusters IV-V) (Fig. 1c, Supplementary Table 6). However, only 830 of the 5,829 gene-associated dATAC regions associate to DEGs (adjusted P≤0.05, log_2_(FC)≥2; Extended Data Fig. 1d; Supplementary Table 5) suggesting that chromatin accessibility is regulated independent of transcriptional control^15–17^. The correlation of chromatin accessibility changes and changes in transcription is exemplified in genome browser views of regulatory sites of ME genes *Gsc*^18^ (cluster I) and *Mesp1*^19, 20^ (cluster II) (Fig. 1d), and anterior epiblast/early NE and pluripotency genes of clusters IV and V, such as *Pou3f1*^21^ and *Phox2b*^22^ (Fig. 1e).

In summary, the temporal mapping of chromatin accessibility changes during ME lineage specification revealed highly dynamic, genome-wide chromatin remodelling, which occurs at gene loci seemingly independent of active transcription at this differentiation stage.

### EOMES binds dynamically opening ME enhancers

The first 24 hrs of early ME differentiation from primed pluripotency (d2-d3) capture the majority of dynamically closing and opening accessible sites (Fig. 1b, Extended Data Fig. 1a). Thus, we focussed the following analyses of dynamic chromatin accessibility changes to this period and grouped all ATAC sites detected between d2-d3 (n=32,392) into closing, stable and opening sites (Fig. 2a, Supplementary Table 7). The analysis of genomic peak distribution shows that opening and closing ATAC sites enrich for enhancer-associated intronic and intergenic regions, while stable ATAC sites are predominantly located at promoter regions (Extended Data Fig. 2d). This genomic distribution of closing/opening and stable sites to enhancers and promoters, respectively, was confirmed by ChIP-seq (chromatin immunoprecipitation followed by deep sequencing) analyses for histone marks H3K4me1^23^ (enhancers), H3K27ac^24^ (active enhancers and promoters), and H3K4me3^25^ (promoters) (Fig. 2b, Extended Data Fig. 2a-c)^26, 27^. GO term enrichment analysis of ATAC-site associated genes revealed neuronal and stem cell programmes enriched at closing sites, primitive streak and ME programmes enriched at opening ATAC sites, and less specific terms, unrelated to germ layer specification enriched at stable ATAC sites (Extended Data Fig. 2e, Supplementary Table 8).

**Fig. 2.**
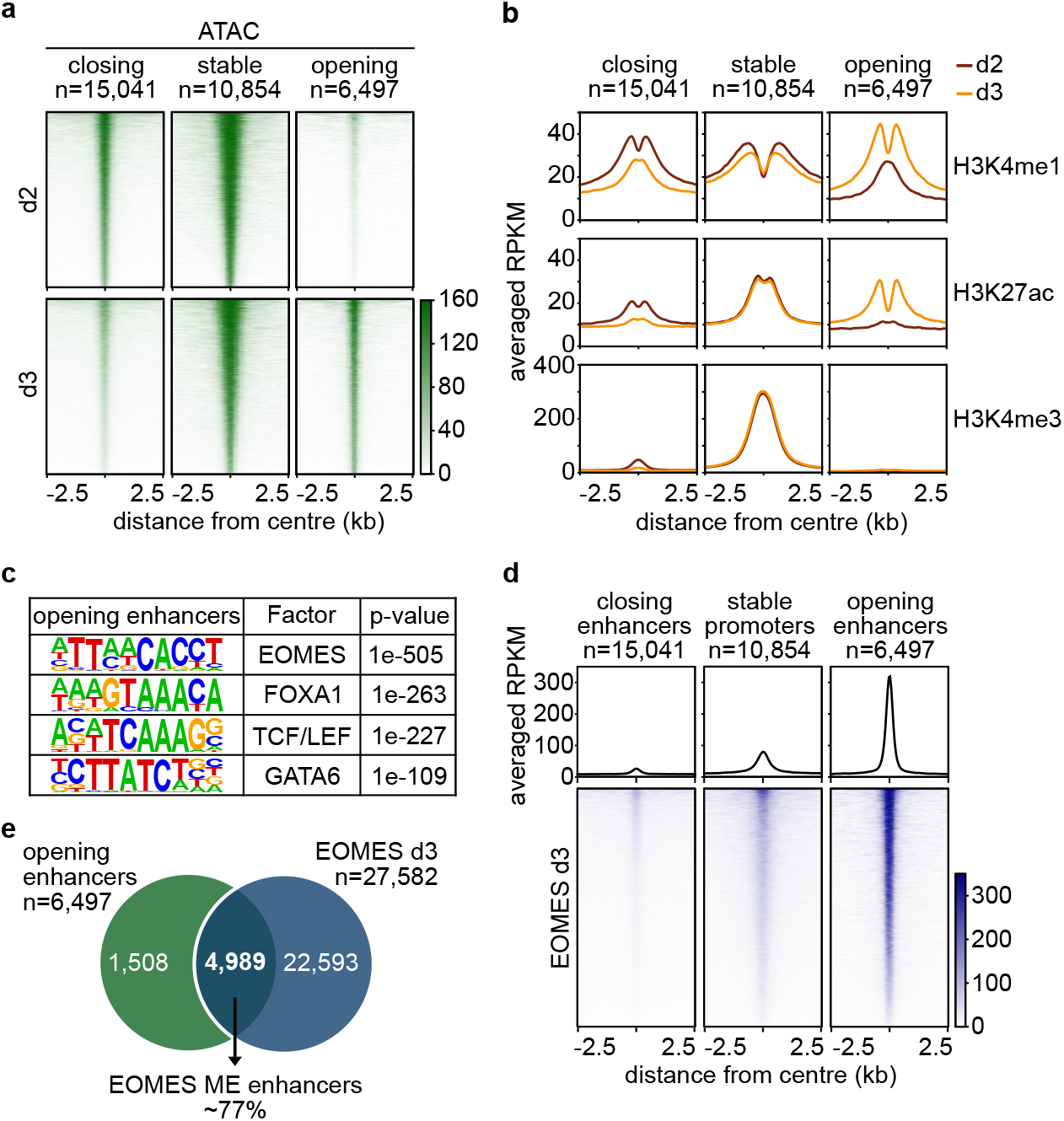
EOMES binds to opening ME enhancers. **a**, Heat maps of ATAC-seq sites ±2.5 kb around peak centre on d2 and d3 of ME differentiation. All ATAC sites were grouped into closing (d2>d3; n=15,041), stable (d2=d3; n=10,682), and opening (d2<d3; n=6,497) regions. **b**, Meta plot representation of ChIP-seq signals for histone modifications H3K4me1, H3K27ac and H3K4me3 centred ±2.5 kb around closing, stable and opening ATAC regions on d2 and d3 of differentiation. Closing and opening regions are enriched for the enhancer marks H3K4me1 and H3K27ac on d2 or d3, respectively. Stable regions show enrichment for the promoter mark H3K4me3. **c**, Transcription factor (TF) binding-motif analysis of n=6,497 opening enhancers using HOMER showing enrichment of motifs for ME TFs. PWMs (positional weight matrices) and p-values for top enriched motifs are indicated. **d**, Meta plots and heat maps of EOMES ChIP-seq enrichment centred ±2.5 kb around the peaks centre of closing, stable and opening ATAC regions at d3 of differentiation showing strong enrichment of EOMES at peak summits of opening enhancers. Counts represent the RPKM normalised read density of two biological replicates. **e**, Intersection of opening enhancers and EOMES ChIP-seq peaks on d3 in WT EBs represented as Venn diagram. ∼77% (n=4,989) of all opening enhancers peaks are bound by EOMES. They are referred to as EOMES ME enhancers and used for further analysis throughout the paper.

In conclusion, dynamically opening ATAC sites represent ME enhancers, and closing sites NE and pluripotency enhancers, confirming the previous notion of a preformed, chromatin accessibility-based lineage bias at enhancers, but not promoter regions towards NE in primed pluripotent cells (d2). Promoter regions are generally less dynamic in respect to analysed histone modifications and chromatin accessibility, which generally remains high during the analysed period from d2-d3 of ME differentiation.

To identify candidate, sequence-specific TFs that control CREs contained in the three classes of ATAC sites (closing NE-related enhancers, opening ME-related enhancers, and stable promoters, Fig. 2a), we performed TF binding motif analysis (Fig. 2c, Extended Data Fig. 3a, b). In *de novo* opening enhancers, we find strong enrichment of motifs of early ME TFs, foremost the motif of the Tbx TF EOMES^8, 28^, and others including FOXA^29, 30^, TCF/LEF^31^ and GATA^32^ TFs (Fig. 2c). Binding motifs enriched in closing enhancers and in stable promoter regions contain those for NE/pluripotency TFs such as SOX2, OCT4, and NANOG (Extended Data Fig. 3a, b), and motifs for less lineage specific TFs such as SP5 (Extended Data Fig. 3b).

The Tbx TF *Eomes* is indispensable for ME lineage differentiation^8, 33–35^, and expression precedes other ME TFs during differentiation (Extended Data Fig. 1c, Extended Data Fig. 3c, Fig. 1c (cluster I)). Accordingly, *Eomes* represents a highly promising candidate for the regulation of early chromatin accessibility changes at ME enhancers. Thus, we analysed binding of EOMES to the three classes of dynamically opening and closing enhancers, as well as stable promoter ATAC sites in d3 WT EBs by ChIP-seq (Fig. 2d). EOMES is highly enriched at opening ME enhancer regions (Fig. 2d), and notably binds to 77% (4,989 of 6,497) of all *de novo* opening enhancer sites (Fig. 2e, Supplementary Table 9). In contrast, EOMES binds to only 5% (702 of 15,041) of closing enhancers sites, and to 14% (1,461 of 10,854) of stable promoter regions (Extended Data Fig. 3d, e). The 4,989 newly established accessible sites that are bound by EOMES were used throughout following analyses of ME lineage specification, and are referred to as EOMES ME enhancers. Expectedly, GO term analysis of EOMES bound ME enhancers showed enrichment for terms associated with ME morphogenesis, such as cardiac differentiation and germ layer formation (Extended Data Fig. 3f).

### *De novo* ME enhancer accessibility relies on EOMES

Next, we investigated if the *de novo* establishment of chromatin accessibility at EOMES ME enhancers also depends on EOMES functions. Here, we used previously described *Eomes*^-/-^ and *Brachyury*^-/-^ double knock-out (dKO) mESCs^8^ (Fig. 3a, b). In these cells the closely related, but later acting Tbx TF *Brachyury* is additionally deleted, since *Brachyury* gains premature functions in, and partially compensates for the absence of *Eomes*^36^. dKO cells fail to specify ME cell lineages^8^ as confirmed by comparative RNA-seq of WT and dKO EBs at d3 (adjusted P≤0.05, log_2_(FC)≥2; Extended Data Fig. 4a, Supplementary Table 10). In dKO EBs, the analysis of chromatin accessibility by ATAC-seq and ChIP-seq of the active histone mark H3K27ac at EOMES ME enhancers showed a drastic loss of enhancer accessibility and activity, respectively (Fig. 3c), accompanied by the gross reduction of the enhancer mark H3K4me1 (Extended Data Fig. 4b, c).

**Fig. 3.**
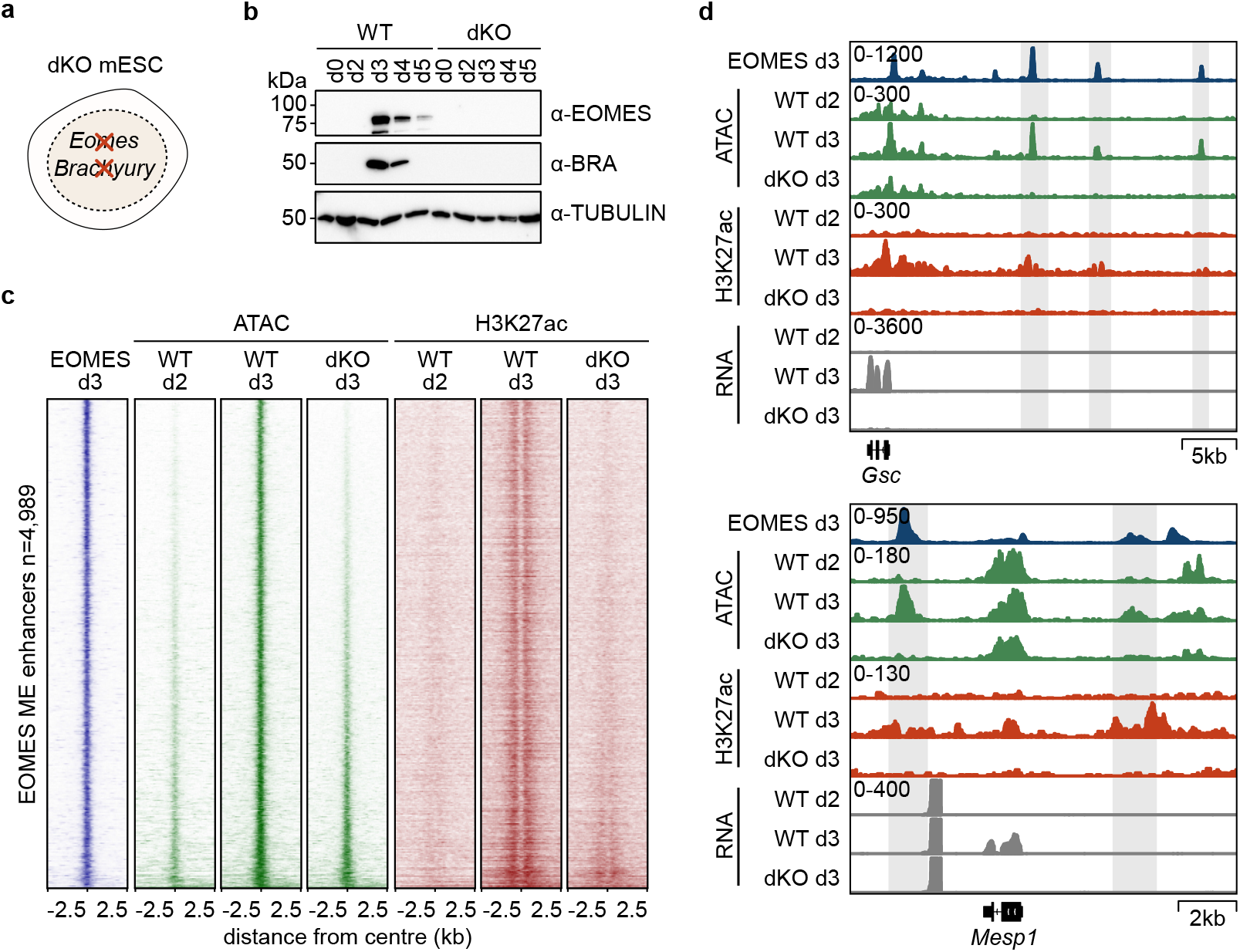
Dynamic opening of ME enhancers relies on EOMES functions. **a**, Schematic of *Eomes* and *Brachyury* double knock-out (dKO) mESCs. **b**, Immunoblots of whole cell lysates from WT and dKO cells during five days of ME differentiation showing the dynamic expression of EOMES and BRACHYURY in WT cells and absence in dKO cells. **c**, Heat maps of EOMES ChIP-seq (blue), ATAC-seq (green) and H3K27ac ChIP-seq (red) in WT and dKO EBs at d2 and d3 of ME differentiation centred ±2.5 kb around n=4,989 EOMES ME enhancers showing significantly reduced ATAC and H3K27ac signals in d3 dKO cells. EOMES ME enhancers are ranked by ascending accessibility in dKO EBs, and the same ranking was used for all heat maps. **d**, Genome browser views of EOMES ChIP-seq (blue), ATAC-seq (green), H3K27ac ChIP-seq (red) and RNA-seq (grey) of EOMES regulated ME target genes *Gsc* and *Mesp1* in WT and dKO differentiated EBs at indicated time points d2 and d3. EOMES binding at opening enhancers is highlighted by grey boxes. Tracks show RPKM normalised counts of at least two merged replicates.

Single track genome browser views of previously described EOMES-dependent target genes *Gsc* and *Mesp1*^33^ further underscore the deficit to gain accessibility at early ME enhancers in dKO cells devoid of Tbx factor activities (Fig. 3d, Extended Data Fig. 4c).

In conclusion, presented data indicate that the *de novo* establishment of chromatin accessibility at ME enhancers that forms within the first 24 hrs after onset of ME specification crucially relies on EOMES, or on compensatory functions by the related Tbx factor BRACHYURY.

### EOMES recruits the canonical SWI/SNF complex

To define the molecular functions of EOMES for the establishment of chromatin accessibility at ME enhancers, we analysed the chromatin-associated protein interactome of EOMES and of the related TF BRACHYURY. We performed ChIP-MS (chromatin immunoprecipitation followed by mass spectrometry)^37^ from dKO EBs expressing C-terminally tagged EOMES:GFP or BRACHYURY:GFP using α-GFP antibody. Doxycycline (dox)-induced expression of GFP-tagged Tbx factors fully rescues endogenous gene functions in dKO cells^8^ (Fig. 4a). Following α-GFP antibody pull-down, the co-precipitating proteins were identified by label-free quantitative MS (Fig. 3b)^38^. Among the most significantly co-enriched proteins following EOMES:GFP pulldown when compared to non-induced (nodox treated) EBs (FDR≤0.05) were multiple subunits of the canonical ATP-dependent chromatin remodelling complex SWI/SNF (SWItch/Sucrose Non-Fermentable) (Fig 4c, d, Supplementary Table 11). SWI/SNF plays crucial roles in changing chromatin accessibility states during development and in disease, including the regulation of early ME genes during lineage specification^39, 40^.

**Fig. 4.**
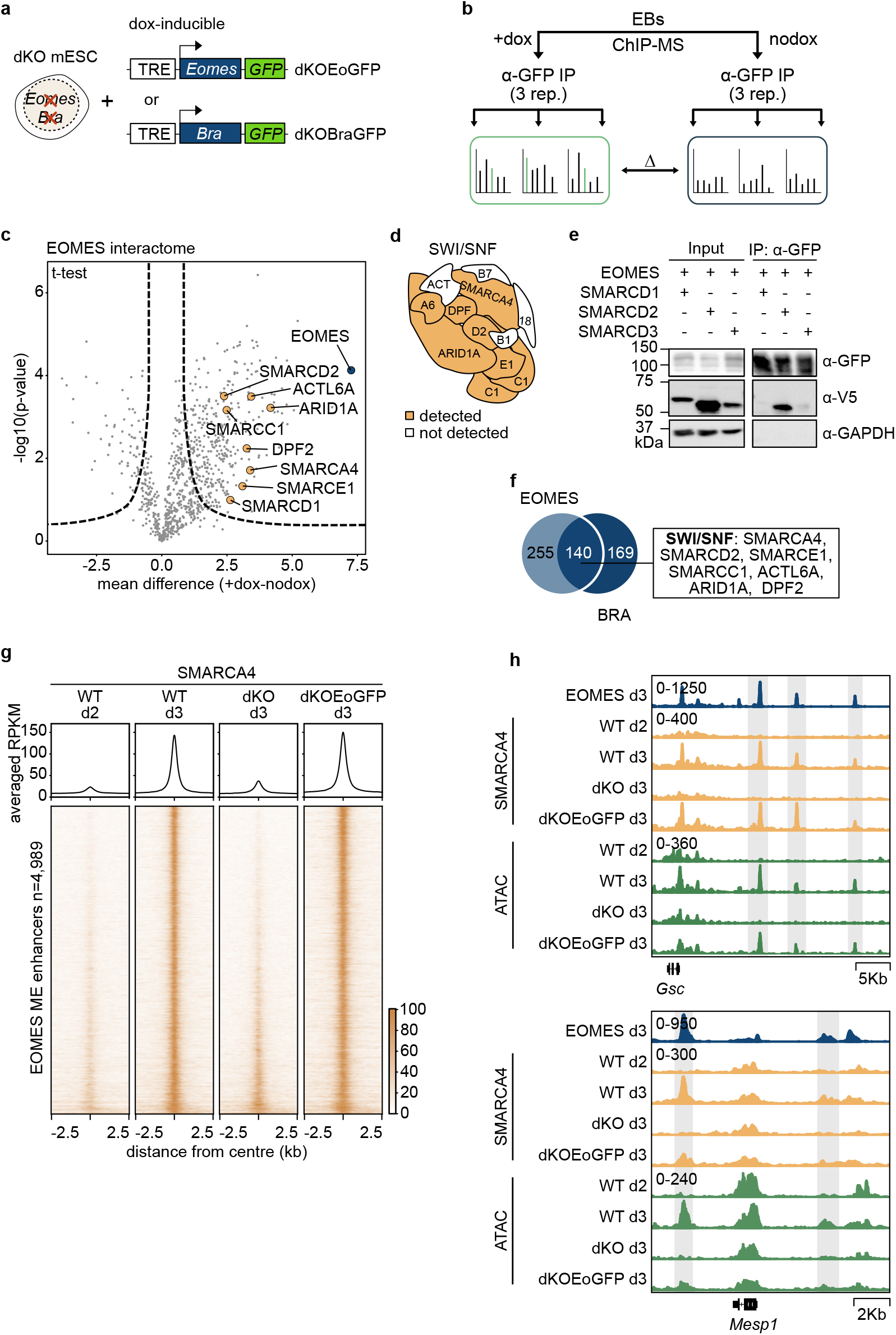
EOMES recruits the canonical SWI/SNF remodelling complex. **a**, Schematic of dKO mESCs harbouring doxycycline (dox)-inducible C-terminally GFP-tagged *Eomes* or *Brachyury* (dKOEoGFP, dKOBraGFP) used for induced expression from d2 of differentiation. **b**, Schematic of quantitative, label-free ChIP-MS (chromatin immunoprecipitation followed by mass spectrometry) of EBs in the presence (+dox) or absence (nodox) of dox-induced expression of GFP-tagged *Eomes* or *Brachyury.* Chromatin was used for α-GFP pull-down and co-precipitated proteins were analysed by MS. Label-free quantification was performed comparing Tbx expressing (+dox) with non-induced (nodox) EBs. **c**, Volcano plot of the EOMES interactome. The negative logarithm of the p-value (two-tailed t-test) on the y-axis is plotted against the difference between the means of logarithmic abundances in +dox treated versus nodox treated cells on the x-axis. Significantly co-precipitated proteins are represented on the right site of the graph (FDR<0.05, s0=1). Subcomponents of the canonical ATP-dependent chromatin remodelling complex SWI/SNF and EOMES are highlighted. **d**, Representation of the full canonical SWI/SNF chromatin remodelling complex^64^. Subunits detected in the EOMES ChIP-MS experiment are highlighted in yellow. C1-SMARCC1, D2-SMARCD2, B1-SMARCB1, E1-SMARCE1, 18-SS18/L1, B7-BCL7A/B/C, ACT-β-Actin, A6-ACTL6A. **e**, Co-IPs of *Eomes:GFP* and *Smarcd1/2/3:V5* by co-transfection in HEK293T cells show binding of SMARCD2 but not SMARCD1 or SMARCD3 to EOMES. α-GAPDH serves as loading control. **f**, Venn diagram showing the overlapping set of 140 co-precipitating proteins detected by ChIP- MS with either EOMES:GFP or BRACHYURY:GFP. Subunits of the SWI/SNF complex detected in both approaches are indicated in the box. **g**, Meta plots and heat maps of SMARCA4 ChIP-seq show the recruitment of SMARCA4 to EOMES ME enhancers (n=4,989) between d2 and d3 in WT cells, which is absent at d3 in dKO cells, and is rescued by the induced expression of EOMES:GFP in dKOEoGFP cells. Regions are ranked according to the reduction of accessibility in dKO cells at d3 (as in Fig. 3c). **h**, Genome browser views of EOMES ChIP-seq (d3, blue), SMARCA4 ChIP-seq (yellow) and ATAC-seq (green) close to ME genes *Gsc* and *Mesp1* in WT at d2 and d3, and dKO and dKOEoGFP EBs at d3 of ME differentiation. SMARCA4 binding and accessibility rely on EOMES-binding to enhancers (grey bar). Tracks show RPKM normalised counts of two merged replicates.

To identify directly EOMES-interacting components we co-expressed V5 epitope-tagged candidate subunits of the canonical SWI/SNF complex together with GFP-tagged EOMES and performed co-IPs. We found co-precipitation of SMARCB1 and SMARCD2, suggesting that these are directly interacting subunits of the SWI/SNF complex (Extended Data Fig. 5a). Notably, from the three highly conserved SMARCD1/2/3 isoforms that only differ in their short N-terminal domain, predominantly SMARCD2 interacts with EOMES (Fig. 4e and Extended Data Fig. 5a). This suggest some degree of specificity for certain SWI/SNF complex subtypes, similar to the specific integration of SMARCD3 to SWI/SNF complexes during cardiac differentiation^41, 42^.

Since EOMES and BRACHYURY share overlapping functions during cell lineage specification^8^, we additionally analysed BRACHYURY in ChIP-MS (Extended Data Fig. 5b, Supplementary Table 12) and found an overlapping set of 140 co-precipitating proteins with EOMES, including the subunits of the SWI/SNF complex (Fig. 3f, Supplementary Table 13). Tagged BRACHYURY co-precipitated the same SWI/SNF subunits SMARCB1 and SMARCD2 as seen with EOMES in co-expression experiments (Extended Data Fig. 5c).

Next, we analysed the dynamics of binding of the SWI/SNF complex to the EOMES ME enhancers that gain accessibility between d2 and d3 (n=4,989; Fig. 2e) using ChIP-seq of SMARCA4 (a.k.a. BRG1) the central ATPase subunit of the canonical SWI/SNF remodelling complex. Expectedly, SMARCA4 is absent from ME enhancers in d2 WT EBs that reflect cells in primed pluripotent state when ME enhancers are inaccessible (Fig. 4g, Fig. 3c). After 24 hrs of differentiation at d3 a total of n=4,989 EOMES ME enhancers become occupied by SMARCA4 and binding is absent in d3 dKO cells (Fig. 4g). However, binding can be fully restored by dox-inducible re-expression of EOMES:GFP showing that site-specific recruitment of SMARCA4 to ME enhancers fully depends on the presence of EOMES, also in the absence of BRACHYURY (Fig. 4g). The EOMES-guided recruitment of SMARCA4 is also shown in track views of known EOMES-regulated enhancer regions of early ME genes *Gsc* and *Mesp1* (Fig. 4h).

In conclusion, EOMES instructs ME lineage commitment by the rapid recruitment of the SWI/SNF chromatin remodelling complex to the broad set of ME enhancers to establish enhancer competence for ME gene programs, as initial regulation to deflect pluripotent cells from their preformed fate-bias towards NE.

### ME enhancer opening is independent of transcription

EOMES is generally considered as a TF controlling target gene expression of cell-type specific gene programs for AM and DE lineages^28, 33–35, 43^. Here, we define EOMES molecular functions as regulator of chromatin for the general initiation of chromatin accessibility at ∼5,000 dynamic ME enhancers, that are associated not only to early AM and DE lineage genes, but also to later transcribed genes (see also Fig. 1b, cluster II genes). Thus, we experimentally tested if the regulation of chromatin accessibility is linked to, and depends on active transcription of enhancer-associated genes.

To control early ME target gene transcription, we reasoned that the expression of the majority of ME lineage-specific genes relies on activities by the Wnt/β-Catenin/Tcf and/or NODAL/SMAD2/3 signalling cascades^9, 44^. Accordingly, we analysed gene expression and chromatin accessibility changes after blocking canonical Wnt and NODAL/SMAD2/3 pathways during onset of early ME program activation (Fig. 5a). Hence, EBs were treated from d2 to d3 of differentiation with the pathway inhibitors SB431542 (SB, blocking NODAL-signalling) and XAV939 (XAV, blocking canonical Wnt-signalling). Simultaneously, EOMES:GFP expression was induced from the dox-inducible locus, bypassing reliance of *Eomes* expression on signalling activities of Wnt/β-Catenin/Tcf and/or NODAL/SMAD2/3^45^ (Fig. 5a).

**Fig. 5.**
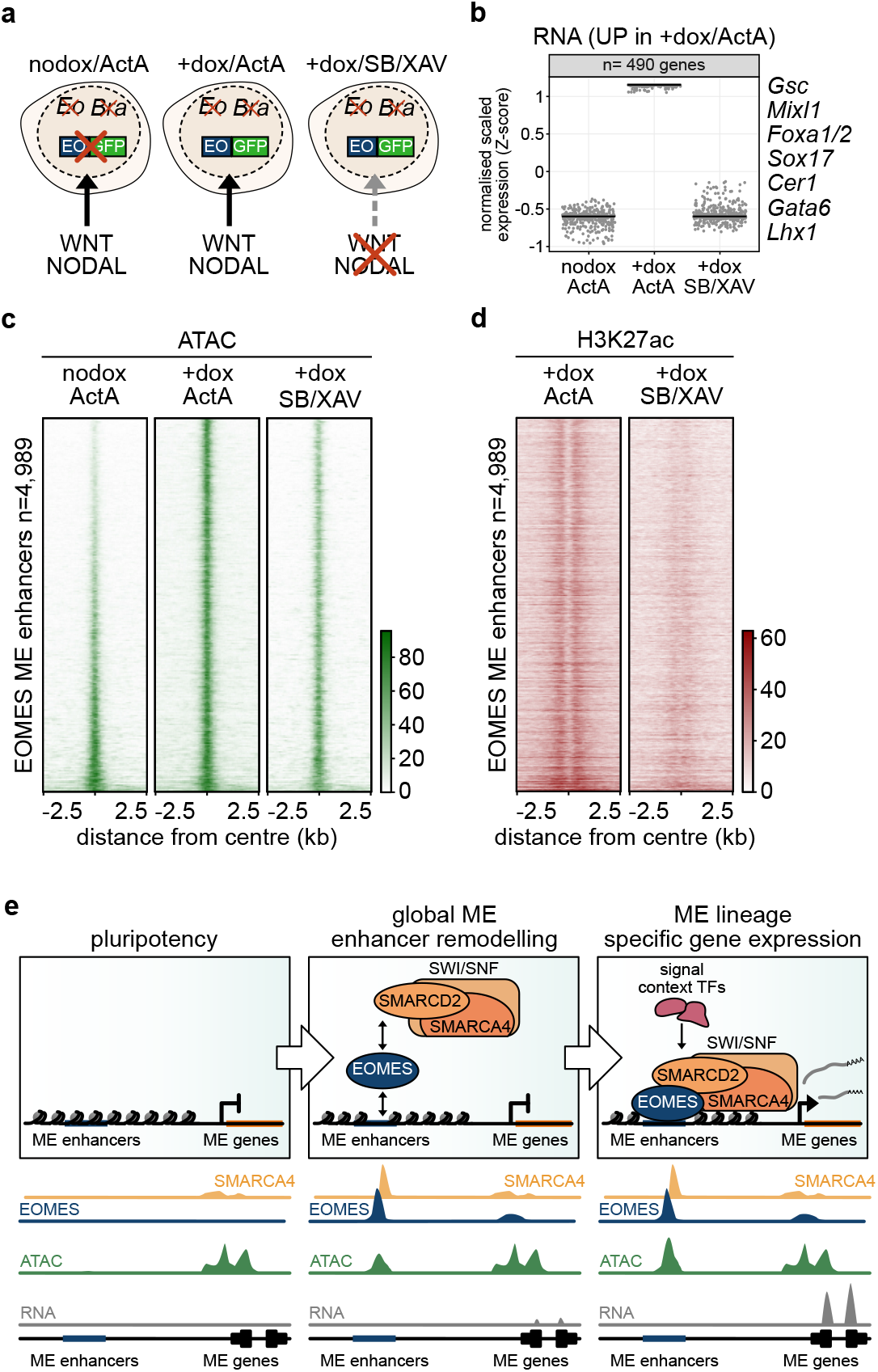
Chromatin accessibility at ME enhancers is established independently of signalling-regulated transcription. **a**, Schematic of EB differentiation of dKOEoGFP cells in 3 conditions: ME induction by Activin A (ActA) in absence (nodox), and presence (+dox) of EOMES:GFP, and in presence (+dox) of EOMES:GFP while blocking signalling by the NODAL/TGFβ inhibitor SB431542 (SB) and the Wnt-inhibitor XAV939 (XAV). Cells were harvested 24 hrs after treatment at d3 for RNA-seq, ATAC-seq, and H3K27ac ChIP-seq. **b**, Dot plots of normalised scaled expression of ME genes that are induced at d3 in dKOEoGFP cells (+dox/ActA) compared to nodox/ActA condition. ME gene expression is absent in nodox/ActA and +dox/SB/XAV treated EBs, indicating the efficient suppression of ME differentiation by blocking NODAL- and Wnt- signalling. Examples for characteristic ME genes of this cluster are indicated. Black lines indicate median values of expression. **c**, Heat maps of ATAC-seq signals in the 3 conditions nodox/ActA, +dox/ActA and +dox/SB/XAV showing that chromatin accessibility is established, however with reduced signal-intensity, at EOMES ME enhancers (n=4,989) in the absence of transcription of associated ME genes when signals are blocked (+dox/SB/XAV). Sites are ranked according to the reduction of accessibility in dKO cells at d3 (as in Fig. 3c) and centred ±2.5 kb around EOMES ME enhancers peaks. **d,** Heat maps of H3K27ac ChIP-seq in +dox/ActA and +dox/SB/XAV condition at d3 ±2.5 kb around EOMES ME enhancers peak centre shows absence of the active enhancer mark in signalling blocking conditions (+dox/SB/XAV). Sites are ranked as in c). **e,** Comprehensive model illustrating the stepwise, EOMES-dependent chromatin remodelling that deflects pluripotent cells from the default NE differentiation path towards initiation of ME lineage commitment. In pluripotent cells the enhancers of ME program genes are in closed chromatin conformation. During gastrulation onset the Tbx TF *Eomes* becomes expressed, globally binds to enhancers of ME program genes and recruits the canonical SWI/SNF chromatin remodelling complex. This recruitment is crucial to generate enhancer competence used by signal-activated transcriptional regulation of ME lineage-specific gene programs.

As expected, SB/XAV treatment abolished ME gene expression, when analysed by RNA-seq (Fig. 5b, Extended Data Fig. 6a; Supplementary Table 14), despite high levels of induced EOMES:GFP (Extended Data Fig. 6b). However, ATAC-seq shows that EOMES ME enhancers nonetheless gain accessibility in SB/XAV conditions, when compared to *Eomes*-deficient, non-induced EBs (Fig. 5c, compare +dox/SB/XAV versus nodox/ActA). Notably, enhancer accessibility in the absence of transcription is less pronounced when compared to EBs that fully engage ME gene expression (Fig. 5c, compare +dox/ActA with +dox/SB/XAV). In contrast to enhancer accessibility the histone mark H3K27ac indicative for active enhancers, fails to be established following SB/XAV treatment (Fig. 5d), suggesting roles of signalling transducers Smads and Tcf/Lef/β-Catenin for enhancer activation, usage and expression control. This is also exemplified in genome browser views of ATAC-seq and H3K27ac tracks of the endoderm marker *Gsc* and mesoderm marker *Mesp1*, that both gain enhancer accessibility in the absence of ME gene transcription by blocking the signalling cascades of Wnt and NODAL/TGFβ (Extended Data Fig. 6c).

In summary, our data show that EOMES regulates ME lineage commitment through establishment of the ME enhancer landscape that generates competence that is used for the activation of signalling-dependent programs of different ME cell types.

## Discussion

This study reveals the molecular regulation that deflects pluripotent cells from the default NE differentiation path towards ME progenitors as illustrated in a comprehensive model (Fig. 5e). We show that the genome-wide competence for transcriptional activation of ME gene programs is initiated by the recruitment of canonical SWI/SNF remodelling complex by the Tbx TF EOMES to establish the accessible ME CREs landscape. This regulation takes place at enhancers, while promoter accessibility is less dynamic. In the absence of Tbx TF activities ME enhancers remain inaccessible and ME gene programs silent. *Eomes* dominates early phases of embryonic ME specification, followed by partially overlapping activities of the related Tbx factor *Brachyury* during formation of more posterior/axial mesoderm derivatives^36^. The Tbx TF- and SWI/SNF- dependent gain of accessibility creates competence for ME gene programs. While chromatin accessibility at ME enhancers is initially established in the absence of transcription (Fig. 5c), it remains to be tested if the persistent presence of EOMES additionally impacts on the efficient transcription of ME target genes. Transcription of lineage specific ME genes relies on activities of signalling pathways, such as Wnt and/or NODAL/TGFβ, and may also depend on the co-binding of additional ME TFs, including FOXA^29^, GATA^32^ and others (Fig. 2c), which fail to become expressed in signalling-inhibitor treated EBs (Fig. 5b). Their binding to ME enhancers might also contribute to robust establishment and maintenance of enhancer accessibility as lineage-defining feature. Additionally, sustained enhancer accessibility as a prerequisite for robust cell lineage identities crucially depends on continuous activities of SWI/SNF chromatin remodelling complex, as previously shown in two studies for the maintenance of pluripotency in ESCs^46, 47^, and for the prevention of trans-differentiation of cardiac cells towards NE cell types^48^. These studies in combination with presented data, demonstrate how the different levels of regulation by chromatin states, TFs and signal-guided transcriptional programmes generate and maintain stable states of lineage-defining gene-programs.

## Methods

### Cell lines

The generation and maintenance of WT (A2lox.Cre^49^), *Eomes* and *Brachyury* double deficient (dKO), and dox-inducible dKOEoGFP and dKOBraGFP mESC lines were previously described^8^. STO feeder cells and HEK293T cells were cultured in DMEM containing 10 % FBS, 1X penicillin/streptomycin and 1X L-glutamine. Cells were grown at 37 °C and 5 % CO_2_.

### EB differentiation to ME

EB formation and differentiation was performed as previously described^8^. In brief, feeder depleted mESCs were seeded for EB formation at 25,000 cells/ml in ESGRO^TM^ Complete Basal Medium (ESGRO, Merck, SF002-500) in non-adhesive 60 mm or 100 mm culture dishes. After 48 hrs of EB formation (d2), differentiation was induced by addition of human recombinant Activin A (ActA, 30 ng/ml, R&D systems, 338-AC-050).

Expression from the Tet-responsible element (TRE) in dKOEoGFP and dKOBraGFP cells was induced by 5 µg/ml doxycycline (dox, Sigma, D9891), NODAL- and Wnt-signalling were blocked by inhibitors SB431542 (10 µM, Tocris, 1614) and XAV939 (2 μM, Tocris, 3748), respectively. Differentiation proceeded up to d4.

### Western Blot

EBs were lysed in ice cold lysis buffer (2% NP40, 20 mM Tris-HCl pH 7.5, 150 mM NaCl, 10% Glycerol) with 1xCPI (Complete protease inhibitors, Roche, 11873580001), 1 mM Na_3_VO_4_, 0.5 mM DTT for 20 min on ice. Whole cell lysates were centrifuged for 15 min at maximum speed at 4°C and protein concentrations measured from the supernatant with a Pierce BCA Protein Assay Kit (Thermo Scientific, 23227). 15 µg protein per lane were resolved on 8% SDS-PAGE, semi-dry blotted on PVDF-membranes and probed for indicated proteins. Primary antibodies used were α-EOMES/TBR2 (ab23345, Abcam, 1:500), α-Brachyury (AF2085, R&D Systems, 1:1,000), α-Brg1 (SMARCA4, ab110641, Abcam, 1:2,500), α-GAPDH (R960-25, Invitrogen, 1:2,500), α-GFP (NB600-308, Novus Biologicals, 1:5,000), α-Tubulin (86298S, Cell Signalling, 1:2,500).

### Chromatin immunoprecipitation followed by deep sequencing (ChIP-seq)

For ChIP-seq the Diagenode iDeal ChIP-seq kit for Transcription Factors (Diagenode, C01010055) was used. In brief, single cell suspensions from EBs were generated by trypsinization, and approximately 10^7^ cells were used per TFs ChIP, and 10^6^ cells were used per histone ChIP. For TFs ChIP chromatin was double-crosslinked for 20 min in 2 mM disuccinimidylglutarate (DSG, Thermo Fisher, 20593) in DMEM, and for additional 10 min supplemented with 1% formaldehyde (Thermo Scientific, 28908). For histone ChIP chromatin was single crosslinked in 1% formaldehyde (Thermo Scientific, 28908) in DMEM for 8 min. Crosslinking was stopped by adding Glycine following the manual of the kit, cells were centrifuged at 500 x g for 5 min at 4°C, and washed twice with ice cold PBS. Cell lysis was performed according to the manual and adapted to cell numbers for TFs or histone ChIPs. The shearing of the chromatin was performed in a Bioruptor Pico (Diagenode) (6 cycles, 30 seconds “ON”, 30 seconds “OFF”). For immunoprecipitation 20 µg of sheared chromatin per TFs ChIP, or 8 µg of sheared chromatin for histone ChIP was incubated with antibody conjugated DiaMag protein A-coated magnetic beads o/n at 4°C. For TFs ChIPs the following antibodies were used:

α-Tbr2/Eomes (ab23345, Abcam), α-Brg1 (Smarca4) (ab110641, Abcam), α-H3K27ac (ab4729, Abcam), α-H3K4me1 (ab8895, Abcam) and α-H3K4me3 (ab4729, Abcam). Beads were washed, chromatin was decrosslinked and DNA was purified as described in the manual. Library preparation was performed using NEBNext Ultra II DNA Library Prep Kit for Illumina (NEB, E7645S) according to the manual of the kit. Size selection was performed to obtain fragment sizes of 300-600 bp using SPRIselect beads (Beckman, B23318) following the manual of the beads. Fragment size of the libraries was validated on a BioAnalyzer 2100 (Agilent) using High sensitivity DNA Kit (Agilent, 5067-4626). Sequencing was performed at Novogene (UK) Company Limited (Cambridge, United Kingdom) on a NovaSeq 6000.

### Assay for transposase accessible chromatin (ATAC-seq)

The ATAC-seq protocol was adapted from a published protocol^50^. In brief, EBs at d2, d2.5, d3 and d3.5 of differentiation were singularized by trypsinization, washed with ice cold PBS and a total number of 50,000 cells were lysed in 50 µl ATAC-seq lysis buffer. Isolated nuclei were mixed with 50 µl of transposition reaction mix containing Tagment DNA Enzyme 1 (Illumina, 20034197). Samples were incubated using a thermo shaker at 37°C, 600 rpm for 30 min during the transposition reaction. Tagmented DNA was purified using the Qiagen MinElute Kit (Qiagen, 28004). Eluted fragments were amplified by PCR with an appropriate number of PCR-cycles (5-8) using Custom NExtera PCR primers^50^ and the NEBNext High-Fidelity 2x PCR Master Mix (New England Biolabs, M0541S). Amplified DNA was purified using Qiagen MinElute Kit (Qiagen, 28004). A size-selection of libraries was performed using SPRIselect beads (Beckman, B23318) for fragments <1,000 bp following the manual. Size distribution of the libraries was determined with a BioAnalyzer 2100 (Agilent) using the High sensitivity DNA Kit (Agilent, 5067-4626). Sequencing was performed at Novogene (UK) Company Limited (Cambridge, United Kingdom) on a NovaSeq 6000.

### RNA-sequencing (RNA-seq)

Total RNA was isolated from approximately 50 differentiated EBs using RNeasy Mini Kit (Qiagen, 74106). RNA integrity was validated using a BioAnalyzer 2100 (Agilent, Agilent RNA 6000 Pico Kit (Agilent, 5067-1513)). Library preparation (Ultra RNA Library Prep Kit) and sequencing was conducted by Novogene (UK) Company Limited (Cambridge, United Kingdom).

### ChIP-seq data analysis

Low-quality reads and adapter reads were trimmed from raw data using Galaxy platform Trim Galore!^51^ (Galaxy Version 0.4.3.1). Sequencing reads were mapped to the mm10 genome using Galaxy platform Bowtie2^52^ (v2.3.4.3+galaxy0) using paired end settings and duplicates were removed (RmDup^53^ (v2.0.1) with default settings). IGV track BigWig files were generated using bamCoverage^54^ (v3.3.2.0.0) with bin size 10 bases and normalisation to RPKM, with paired-end extension of the fragments. BigWig files of biological duplicates were combined using bigwigCompare^54^ (v3.3.2.0.0) by computing the mean of the signal. Heat maps were generated with ChIP-seq RPKM normalised counts depicted as the intensity of the colour scale, the meta plot was created with averaged RPKM values using deepTools computeMatrix and plotHeatmap (v3.3.2.0.0 and v3.3.2.0.1, respectively^54^) with an mm10 blacklist.

The enriched regions of EOMES ChIP-seq were detected using MACS2^55^ (v2.1.1.20160309.6) with pooled treatment files from two biological replicates using the BAMPE format as an input file and DSGInput BAM file (recently published available under GSM3676129) as control with default settings for building the shifting model: confidence enrichment ratio against background 5-50; band width 300 and q-value of 0.01. High confidence peak list for EOMES ChIP-seq was generated by sorting peaks in tabular format file for pileup values >25 using Filter data on any column using simple expressions (Galaxy Version 1.1.1).

### ATAC-seq data analysis

Processing of ATAC sequencing reads was performed as described for ChIP-seq samples. BAM files and BigWig files were generated as described for ChIP-seq analysis. Peaks were detected using MACS2 (v2.1.1.20160309.6, with BAMPE format of the input file, default settings for building the shifting model (confidence enrichment ratio against background 5-50; band width 300) and minimum q-value cut-off for detection of 0.01.

Differential chromatin accessibility was determined using the R library DiffBind^56^ (v3.6.0). A mm10 blacklist was applied to the analysis, leaving 37,846 unified peak sites (filtered for FDR≤0.05) from consecutive time comparisons between d2-d3.5 of differentiation. Opening, closing and stable ATAC sites between different time point comparisons were defined by filtering DiffBind output file for Fold >1 for opening ATAC, Fold <-1 for closing ATAC and stable ATAC sites were defined by Fold ≥-1 and ≤1 and FDR threshold 0.05. In the DiffBind tool, Fold is defined as: the difference in mean of log2 normalised ATAC read counts. We defined a region as dynamic if this region shows at least once opening or closing behaviour among consecutive time comparisons, otherwise a region is defined as stable.

An informative subset of 10,000 accessible peak sites (dATAC) as given by the default DiffBind parameters were chosen. These 10,000 peaks were then annotated to the nearest gene for further analysis using the R library GREAT^57^ (v1.28.0) using the “oneClosest” rule parameter and a “oneDistance” value of 10 kb to restrict over-annotation of ATAC sites to multiple genes. Multiple dATAC peaks associated to the same gene were merged, producing a unique gene table of 5,829 dATAC associated genes with DiffBind scores per time point corresponding to the normalised read counts. The merged scores were scaled to zero-mean and unit variance, where K-means (k=5) clustering was then performed, using a burnout value of 1,000. The clustered data was visualised via the pheatmap::pheatmap^58^ function (pheatmap v.1.0.10) without using the intrinsic clustering settings.

EOMES bound ME enhancer regions were defined by the intersection of EOMES high confidence peak list and opening ATAC region (d2 vs d3) list using Bedtools Intersect intervals (v2.29.2) with -wa and -u settings (minimal overlap of at least 1bp).

### RNA-seq data analysis

RNA-seq data analysis was performed as previously described^8^. BigWig tracks were generated as described for ChIP-seq data analysis from merged BAM files of replicates using Merge BAM Files (v1.2.0). Differentially expression analysis was carried out using R library DESeq2 (v1.34.0). Briefly, the count data was fitted with a Negative Binomial GLM fitting and Wald statistics were performed on the data, with subsequent filtering for adjusted P≤0.05 and log_2_(FC) as indicated in the figure legends. Prior to fitting and statistical analysis, counts were normalised by library size (DESeq2::estimateSizeFactors) and gene-wise dispersion (DESeq2::estimateDispersions) to normalise the gene counts by gene-wise geometric mean over samples. To reduce background noise in the dKOEoGFP nodox/ActA, +dox/ActA and +dox/SB/XAV analysis, genes were filtered for expression levels >50 normalised counts.The normalised count data was scaled to zero-mean and unit variance and K-means (k=5) clustering was applied using DEGs given by DESeq2. Heatmap visualisation was performed as described in ATAC-seq data analysis section.

To visualise the RNA expression over time at different dATAC clusters DEGs from DESeq2 were intersected with dATAC associated genes. Normalised and Z-score scaled RNA gene expression values of genes within the dATAC clusters were plotted over time. Box and violin plots were generated using the R library ggplot2 (v3.3.6).

### Genomic regions enrichment and Gene Ontology overrepresentation

Genes associated to opening, stable and closing ATAC sites or EOMES ME enhancers were determined using Genome Regions Enrichment Annotations Tool (GREAT, v4.0.4^57^) with basal plus extension settings. Gene Ontology analysis results from GREAT were used to represent enriched terms at these regions.

### Genomic peak distribution

For visualisation and annotation of genomic peaks of closing, stable and opening ATAC regions ChIPseeker^59^ tool was performed with default settings in Galaxy (Version 1.18.0+galaxy1).

### Motif enrichment analysis

Motif enrichment analysis was performed in Galaxy using findMotifsGenome (HOMER)^60^ (Galaxy Version 4.11+galaxy2) with installed genome mm10 Full source FASTA sequence on peak summits +/- 150 bp based on DiffBind output peak files for closing, stable or opening ATAC peaks.

### Chromatin Immunoprecipitation-Mass spectrometry (ChIP-MS)

ChIP-MS of cross-linked chromatin was performed as described previously^37^ with some modifications adapted for mESCs. In brief, dKOEoGFP or dKOBraGFP mESCs (25,000 cells/ml) were grown as EBs in serum free medium and treated with ActA (12.5 ng/ml) with or without dox (5 µg/ml) from d2 until d4. Approximately 30-50 million cells per condition (+dox or nodox) were crosslinked with 1% formaldehyde (Fisher Scientific, cat. no. 28906) for 8 min at RT. 10 µl α-GFP antibody (ab290, Abcam) pre-bound to 100 µl SureBeads^TM^ Protein G beads (BioRad, 1614821) was used per IP. Immunoprecipitated proteins were washed and subject to an on-bead trypsin digest.

### Sample preparation for MS

The tryptic digest was desalted using HyperSep^TM^ SpinTips (Thermo Scientific) according to the instructions with some modifications. Binding solution contained 0.1% TFA and releasing solution contained 65% acetonitrile (ACN) in ddH_2_O. HyperSep^TM^ SpinTips were prepared according to the manual prior to sample loading, repeated washes and elution in 50 µl of releasing solution. Eluates were lyophilised in a vacuum concentrator (45°C, 600 rpm, -15 mbar vacuum, 20 mbar safety pressure). Samples were stored at -80°C prior to LC-MS/MS analysis.

### MS measurements

Lyophilised peptides were analysed by an Orbitrap Q-Exactive Plus (Thermo Scientific) mass spectrometer coupled to an EASY nano-LC 1000 (Thermo Scientific) with a flow rate of 300 nl/min. Samples were mounted on a C18 separating column (2 µm particle size, 100 Å pore size, length 150 mm, inner diameter 50 µm, Thermo Fisher Scientific, 164711). An acetonitrile gradient (0–80%) was applied for reverse phase chromatography. A maximum of 10 MS/MS scans following each MS1 scan was selected in a data dependent acquisition (DDA) mode.

### MS data analysis

For peptide identification, raw MS spectra were loaded into MaxQuant^61^ (v1.5.2.8) and analysed against the Uniprot mouse annotated protein database (June 2017). A decoy database was created by using the reverts function. Fixed modifications for peptide-spectrum matching included carbamidomethyl cysteine and variable modifications of methionine oxidation and N-terminal acetylation. Missed cleavages by trypsin were set to zero and a protein was identified by at least one peptide. The false discovery rate (FDR) at both peptide and protein level was set to 0.01. MaxLFQ algorithm was selected to quantify the label values. Selection for significant enriched proteins was performed in Perseus^62, 63^ (v1.5.0.0). Perseus was used to filter contaminants and reverse hits. Label-free quantification values were transformed to log2 and grouped according to sample condition (nodox or +dox). Proteins measured in at least three times in one condition (nodox or +dox) were accepted. Proteins were visualised in Perseus in volcano plots, therefore missing values were replaced by values from the normal distribution. Significance was calculated by applying two-tailed Students t-test with FDR threshold 0.05 and s0=1.

### Quantification and statistical analysis

All ChIP- and ATAC-seq experiments were performed as two biological independent experiments, all RNA-seq experiments as 3-4 biological independent experiments. All ChIP-MS experiments were performed from three biological independent experiments.

### Co-immunoprecipitation in HEK293T cells

HEK293T cells were transiently co-transfected with either 10 µg GFP-tagged EOMES or BRACHYURY pcDNA6 expression vector and 10 µg V5-tagged SWI/SNF subcomponents pcDNA6 expression vector. One day after transfection cells were lysed on the plate with 1 ml of IP lysis buffer (0.1% Tergitol-type NP40, 50 mM Tris HCL pH 8.0, 300 mM NaCl, 10 mM MgCl_2_, 10% Glycerol) complemented with 1x CPI (Complete protease inhibitors, Roche, 11873580001), 1 mM Na_3_VO_4_ and 0.5 mM DTT for 30 min under constant agitation at 4°C. Whole-cell lysate was collected and the samples were centrifuged for 20 min, 12,000 rpm at 4°C. Whole cell lysate was incubated with 1 µl α-GFP antibody (NB600-308, Novus Biologicals) coupled ProteinA Sepharose beads (VWR, GE17-0780-01) overnight under constant rotation at 4°C. After serial washes in IP lysis buffer the samples were denaturized in 1x sample buffer. Total IP or 15 µg of input samples were resolved by SDS-PAGE and subjected to immunoblot analysis. The membrane was probed for α-GFP (NB600-308, Novus Biologicals, 1:5,000), α-V5 (R960-25, Invitrogen, 1:2,500) and α-GAPDH (R960-25, Invitrogen, 1:2,500).

### Data availability

RNA-seq, ATAC-seq, and ChIP-seq data of differentiated mESCs have been deposited in the Gene Expression Omnibus (GEO) under accession code GSE218839. Input data for peak calling with MACS2 was previously published and is accessible under GSM3676129. The mass spectrometry proteomics data have been deposited to the ProteomeXchange Consortium (http://proteomecentral.proteomexchange.org) with the dataset identifier PXD038355.

### Code availability

All customised computational code used in this study are available under https://gitlab.com/mtekman/eomes-brachyury-analysis/.

## Acknowledgements

This work is dedicated to Hans-Henning Arnold (†12.12.2022), a pioneer in myogenic lineage specification. We thank T. Bass and S. Koidl for technical assistance, M. Kyba for the A2lox.Cre mESC line, the Freiburg Galaxy Team, in particular Björn Grüning, Bioinformatics, University of Freiburg (Germany) funded by the Collaborative Research Centre 992 Medical Epigenetics (DFG grant SFB 992/1 2012) and the German Federal Ministry of Education and Research BMBF grant 031 A538A de.NBI-RBC, the BIOSS Toolbox Cloning Service (Freiburg, Germany), and G. Walz (University clinic, Freiburg, Germany) for providing plasmids. This study was supported by the German Research Foundation (DFG) through the Heisenberg Program (AR 732/3-1), project grant (AR 732/2-1), project P7 of CRC 1453 (project ID 43198400), project A03 of CRC 850 (project ID 89986987), project A08 of CRC 992 (project ID 192904750) to S.J.A., DDG project grants 446058856, 466359513, 444936968, 405351425, 431336276, 43198400 (SFB 1453 “NephGen”), 441891347 (SFB 1479 “OncoEscape”), 423813989 (GRK 2606 “ProtPath”), and 322977937 (GRK 2344 “MeInBio”) to O.S.; project A07 of CRC 992 (project ID 192904750) to H.T.M.T; and Germany’s Excellence Strategy (CIBSS – EXC-2189 – Project ID 390939984) to S.J.A. and S.P.; K.M.S is funded by the EQUIP Program for Medical Scientist, Faculty of Medicine, University Freiburg, and by a CRC 992 MEDEP-Fellowship.

## Author contributions

Project design and direction: C.M.S. and S.J.A. mESCs engineering, in vitro differentiation, RNA-seq, ChIP-seq and data analysis: C.M.S. and S.-L.M. ATAC-seq: L.Z. Experimental design Co-IPs and proteomics: O.S. and H.T.M.T. mass spectrometry and data analysis: C.M.S. and O.S. Western blot analysis: C.M.S and S.-L.M. Bioinformatic data analysis: C.M.S. and M.T. Additional RNA-seq, ATAC-seq and ChIP-seq data analysis S.P., K.M.S., S.P. and H.T.M.T. Figures, manuscript writing: C.M.S., S.P. and S.J.A., with input from all authors.

---

**Extended Data Fig. 1.**
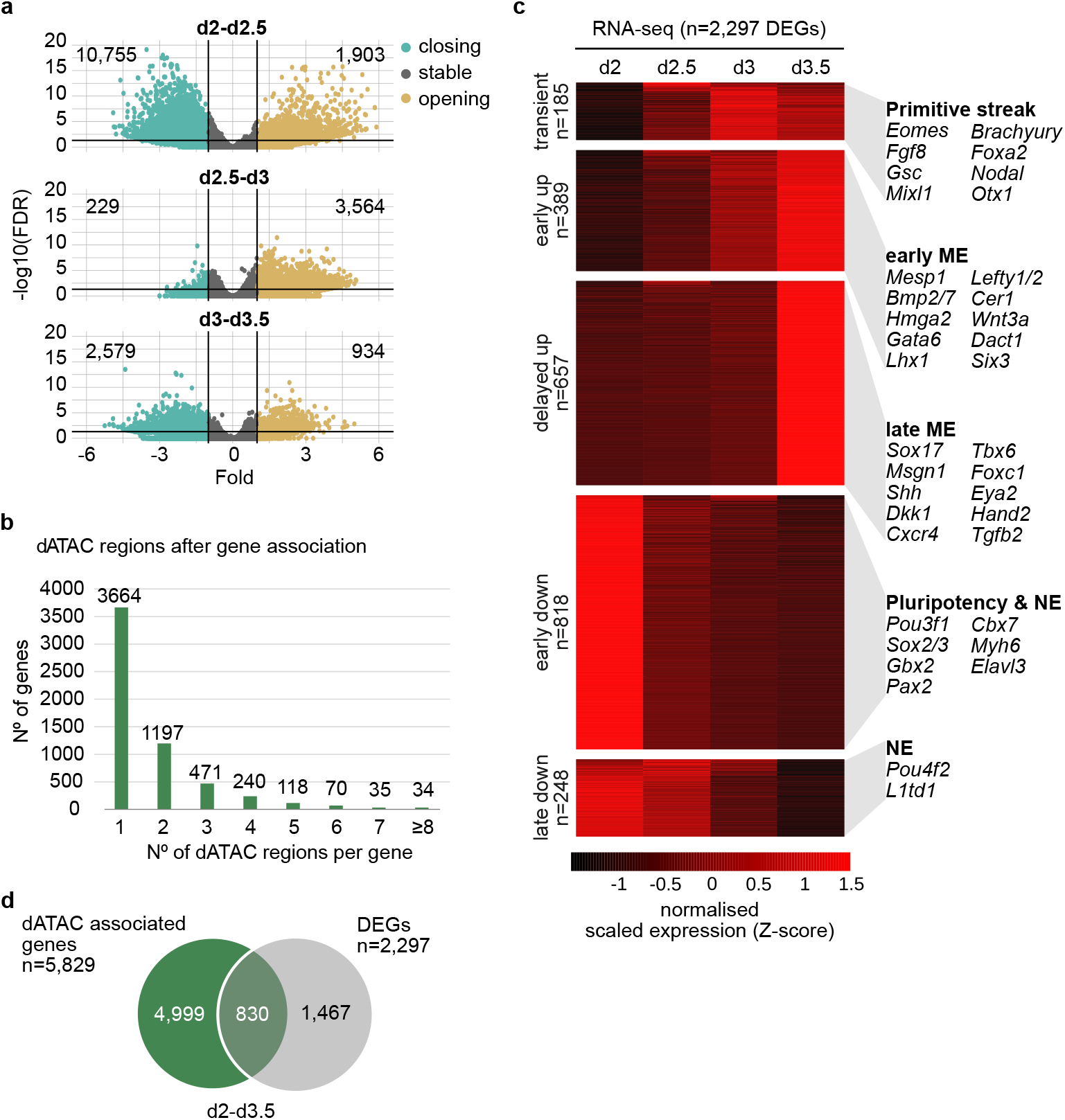
Association of dynamic chromatin accessibility with gene expression changes. **a**, Volcano plots showing dynamics of chromatin accessibility changes between indicated time points. The x-axis indicates Fold differences at ATAC sites and the y-axis the corresponding -log10(FDR). The numbers of closing and opening ATAC sites between time points are indicated. Thresholds for FDR of 0.05 and Fold of -1 and 1 are indicated by lines. **b**, Histogram showing the frequencies of dATAC sites associated to the nearest single. The majority of the all 5,829 genes associate to a single dATAC site (3,664). **c**, Heat map of differentially expressed genes (DEGs; adjusted P≤0.05, log_2_(FC)≥2) by RNA-seq from WT EBs differentiated to ME from d 2 tod3.5 in 12 hrs intervals. DEGs were grouped into 5 clusters by K-mean (k=5). Representative marker genes for different cell types within each cluster are indicated. Significance was assessed by DEseq2 on the basis of two-sided Wald test with Benjamini-Hochberg adjusted *P* values. **d**, Venn-diagram of the intersection of dATAC associated genes with DEGs between d2 and d3.5 demonstrating that of 5,829 dATAC associated genes only 830 are differentially expressed (DEGs).

**Extended Data Fig. 2.**
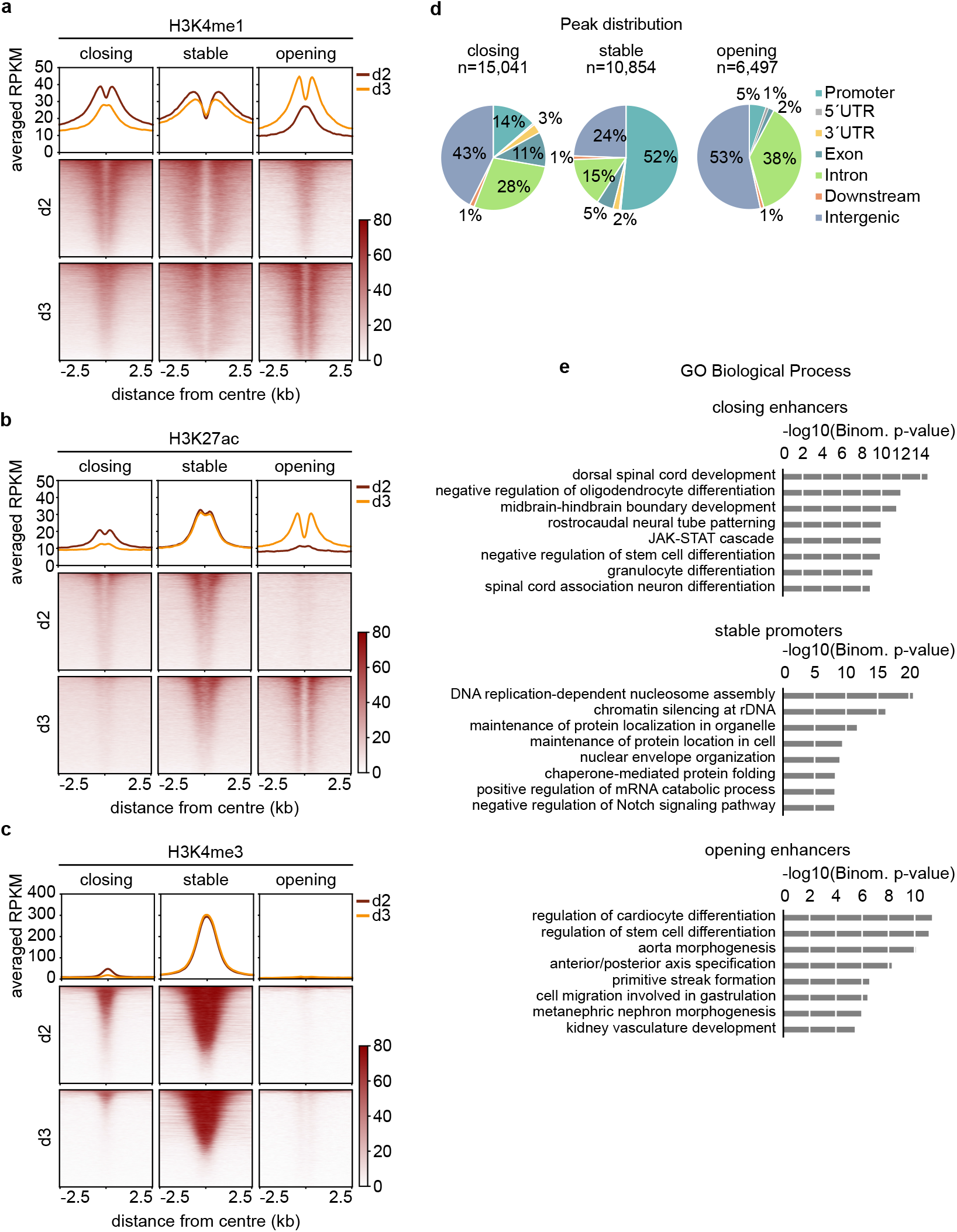
Characterisation of closing, stable and opening ATAC sites. **a-c**, Meta plots and heat maps of histone modifications (H3K4me1, H3K27ac and H3K4me3) at closing, stable and opening ATAC regions at d2 and d3 of ME differentiation. ChIP-seq enrichments are centred ±2.5 kb around peak summit of ATAC sites. **d**, Pie charts showing the genomic peak distribution of closing, stable and opening ATAC sites. Closing and opening regions are located predominantly within intronic and intergenic regions, while stable regions are enriched at promoters. **e**, Gene ontology (GO) analysis for Biological Process of closing, stable and opening ATAC associated genes. For closing regions, there is an enrichment for neuronal terms. Stable sites are enriched for general cellular functions. Opening sites are enriched for terms associated with ME development.

**Extended Data Fig. 3.**
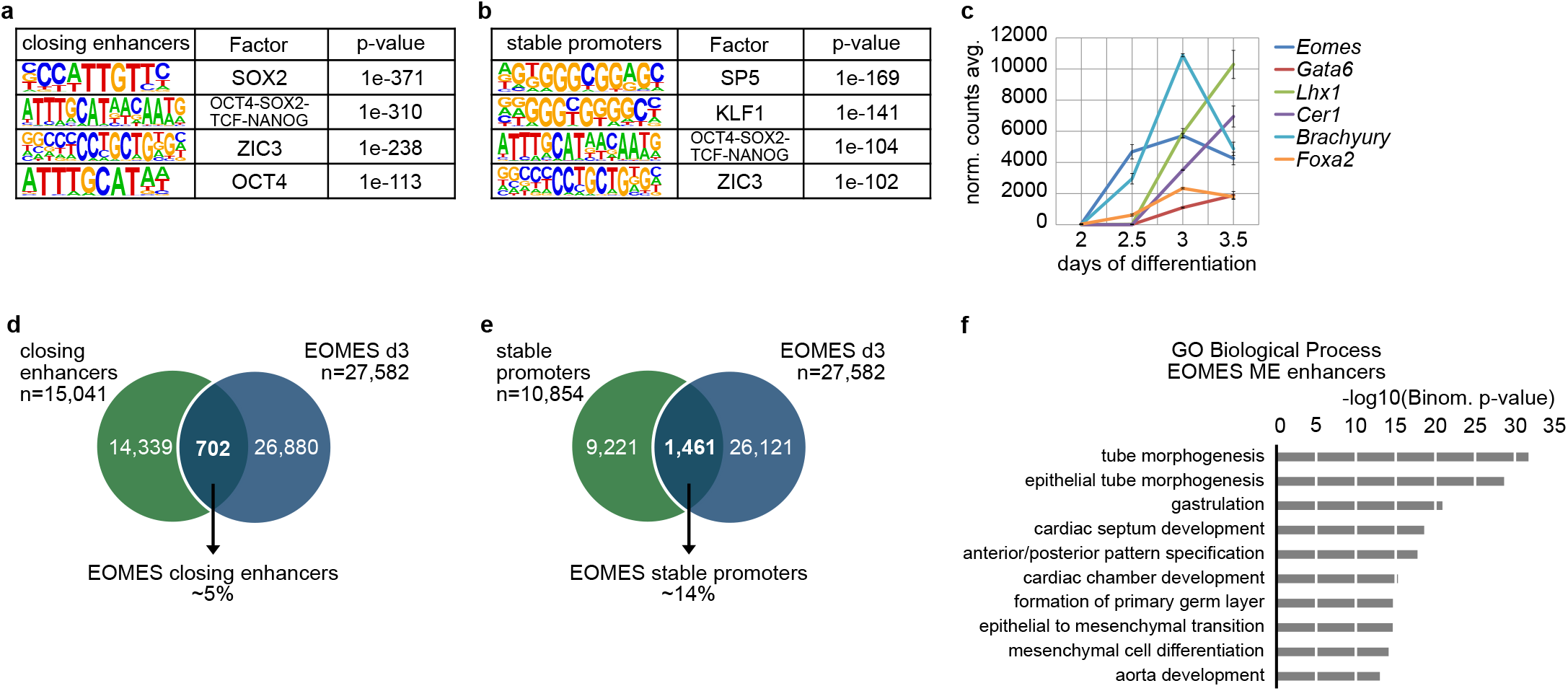
Characterisation of closing enhancers and stable promoters. **a, b**, Tables of enriched TF motifs by HOMER analysis showing the most enriched TF binding motifs as PWM (position weight matrix) within closing and stable ATAC sites during early ME differentiation (d2 – d3), corresponding TFs are indicated and p-values shown. **c,** RNA-expression of indicated, early ME genes during the course of early ME differentiation from d2 to d3.5. Expression values are shown as averaged, normalised counts of RNA-seq data. Error bars indicate the standard deviation. **d**, **e**, Venn diagrams showing the intersection of closing or stable ATAC regions with EOMES-binding by ChIP-seq on d3 of WT EBs. Only ∼5% of closing enhancers, and ∼14% of stable promoters are bound by EOMES at d3 of ME differentiation. **f**, GO term analysis of EOMES ME enhancer associated genes using GREAT.

**Extended Data Fig. 4.**
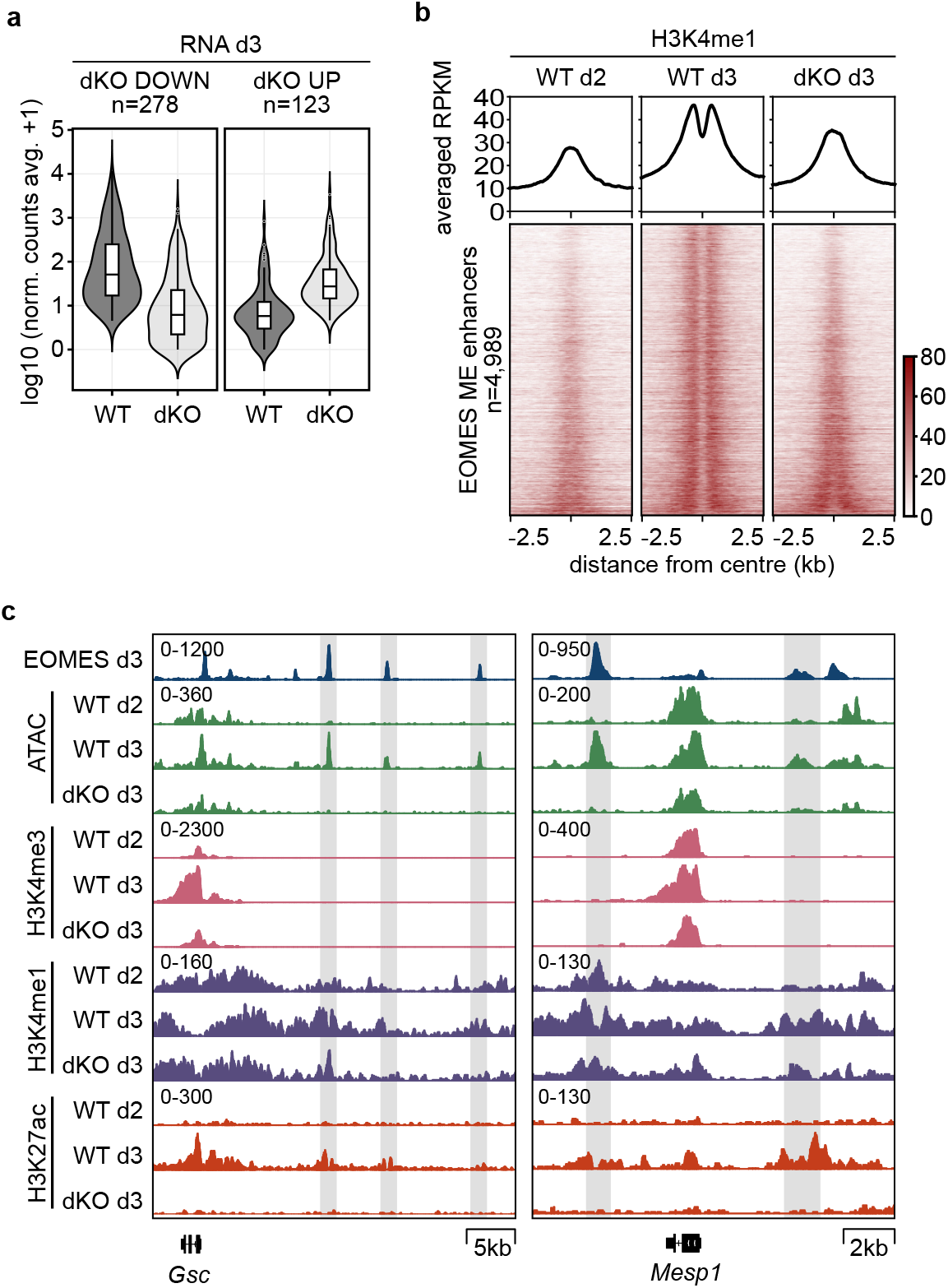
Establishment of EOMES ME enhancers during differentiation. **a**, Violin plots showing the global differences in gene expression between WT and dKO EBs at d3 of ME differentiation. 278 differentially expressed genes (DEGs) are downregulated, and 123 DEGs are upregulated in dKO compared to WT EBs. The y-axis depicts the expression as averaged normalised counts of DEGs in WT vs. dKO EBs (adjusted P≤0.05, log_2_(FC)≥2). Boxes indicate the 25th and 75th percentiles, the median value is indicated by the black line. Whiskers extending no more than 1.5 times the inter-quartile range (IQR). Outliers are indicated by circles. **b,** Meta plots and heat maps of H3K4me1 ChIP-seq enrichment centred ± 2.5 kb around EOMES ME enhancer sites in WT EBs at d2 and d3, and in dKO EBs at d3, showing the establishment of the enhancer mark H3K4me1, sparing the EOMES binding site in WT but not in dKO cells at d3. Sites are ranked according to the reduction of accessibility in dKO cells at d3 (as in Fig. 3c). **c,** Genome browser views at EOMES target genes *Gsc* and *Mesp1*. Shown tracks are EOMES ChIP-seq d3 (blue), ATAC-seq in WT and dKO (green), H3K4me3 (pink, promoter mark), H3K4me1 (purple, enhancer mark), and H3K27ac (red, active promoters and enhancers) ChIP-seq in WT and dKO at d2 and d3 of ME differentiation. Tracks show normalised counts to RPKM of two merged replicates.

**Extended Data Fig. 5.**
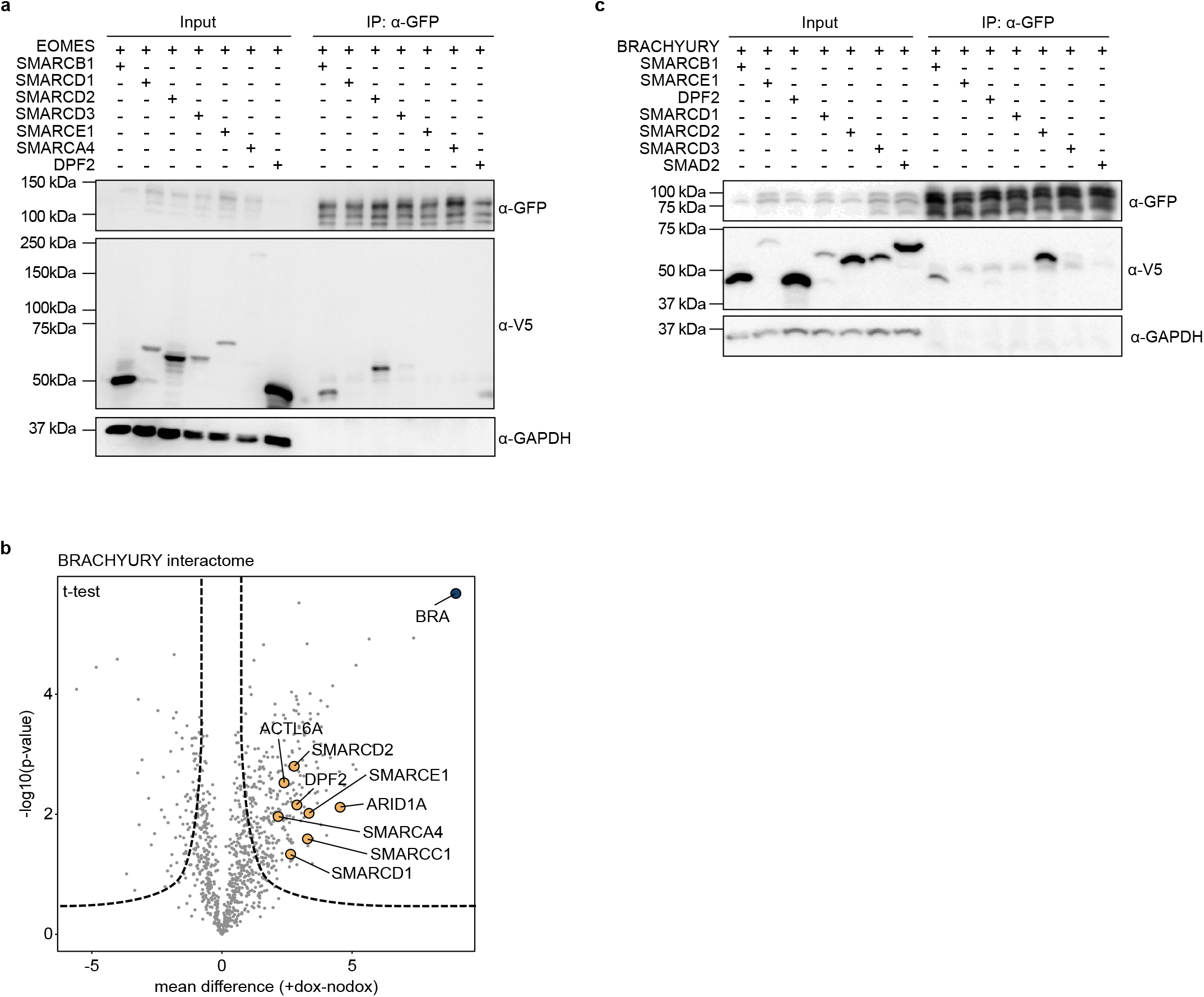
EOMES and BRACHYURY interact with the SWI/SNF chromatin remodelling complex. **a**, Co-IPs of *Eomes:GFP* with *V5-*tagged SWI/SNF candidate subunits after co-transfection in HEK293T cells. IPs from whole cell lysates by α-GFP antibody analysed by immunoblotting and probing for α-GFP and α-V5 antibodies. α-GAPDH serves as control. 15 µg of whole cell lysates served as input controls. SWI/SNF subunits SMARCB1 and SMARCD2 co-precipitate with EOMES. **b**, Volcano plot showing the protein interactome of BRACHYURY following ChIP-MS in differentiated dKOBraGFP EBs. The negative logarithm of the p-value (two-tailed t-test) on the y-axis is plotted against the difference between the means of logarithmic abundances in +dox treated versus nodox treated cells on the x-axis. Significantly co-precipitating proteins are represented on the right site of the graph (FDR<0.05, s0=1). Subcomponents of the canonical ATP-dependent chromatin remodelling complex SWI/SNF are highlighted as yellow circles, and BRACHYURY as blue circle. **c**, Co-IPs of *Brachyury:GFP* and *V5-*tagged SWI/SNF subunits by co-transfection in HEK293T cells. The immunoblot was stained for α-GFP, α-V5 and α-GAPDH. BRACHYURY co-precipitates SMARCB1 and SMARCD2, similar to EOMES as in **a**.

**Extended Data Fig. 6.**
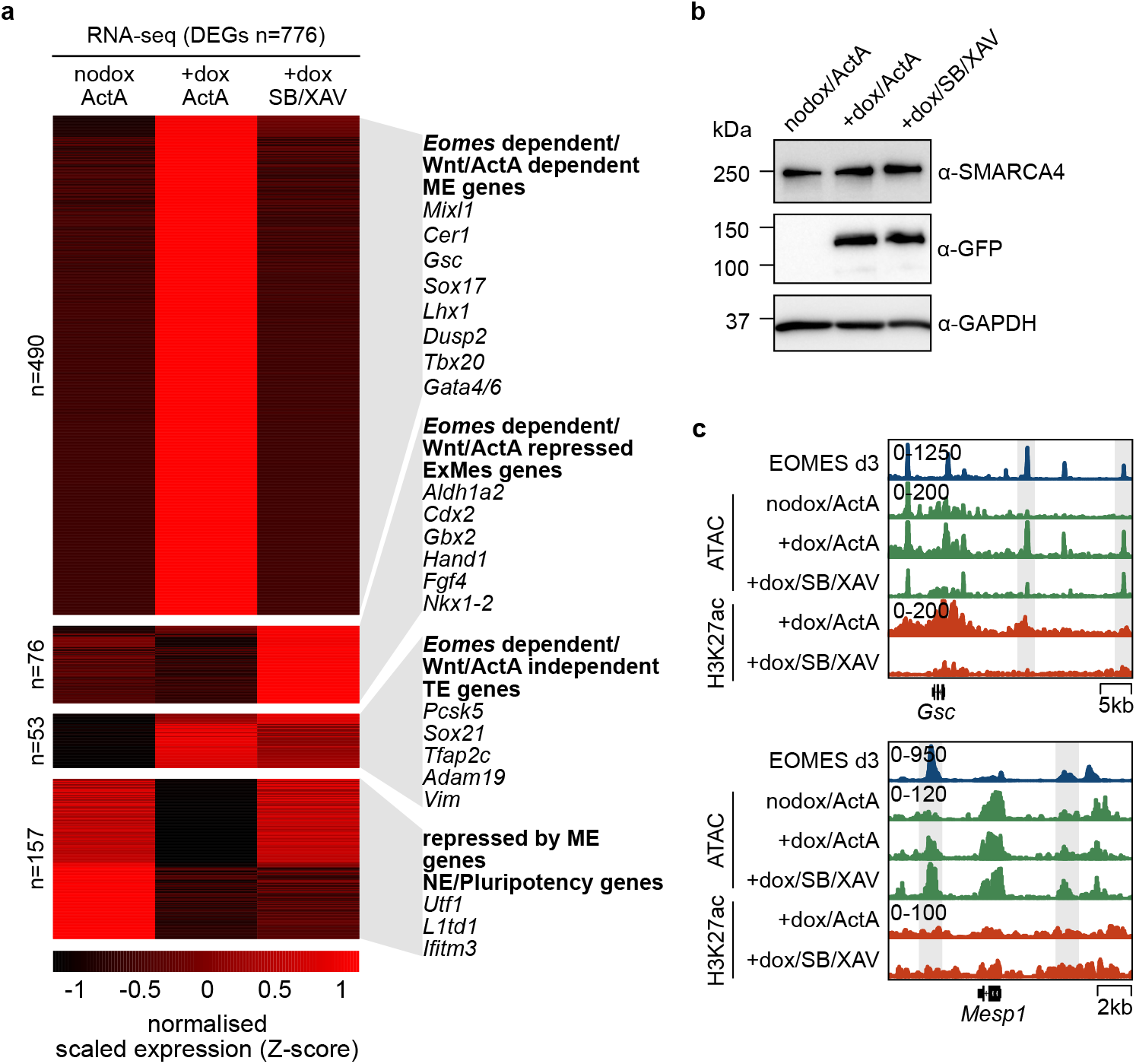
Inhibition of NODAL/TGFβ- and Wnt-signalling abolishes ME gene transcription. a, Heat map of differentially expressed genes (DEGs) of dKOEoGFP EBs at d3 of ME differentiation in conditions of ActA stimulation and inhibition of NODAL/TGFβ - and Wnt-signalling. DEGs (n=776, adjusted P≤0.05, log_2_ (FC)≥2) were clustered by K-means. Representative markers of different gene clusters and their mode of regulation are indicated. Significance was assessed by DEseq2 on the basis of two-sided Wald test with Benjamini-Hochberg adjusted P values. ME - Mesoderm and Endoderm; ExMes - extraembryonic Mesoderm; TE - Trophectoderm; NE - Neuroectoderm. **b,** Immunoblots of whole cell lysates of dKOEoGFP EBs at d3 showing similar levels of induced (+dox) EOMES:GFP expression in Activin A treated (ActA) cells, compared to EBs during inhibition of NODAL/TGFβ- and Wnt-signalling (SB/XAV). In uninduced cells EOMES:GFP fails to be detected. GAPDH serves as loading control. **c**, Exemplary genome browser views of EOMES ChIP-seq (d3 in WT, blue), ATAC-seq (green) and H3K27ac ChIP-seq (red) in dKOEoGFP cells in 3 indicated conditions. The analysis shows increasing enhancer accessibility at ME genes *Gsc* and *Mesp1* at sites of EOMES binding in the absence of transcription in signalling blocking conditions (+dox/SB/XAV). In contrast, H3K27ac fails to be established in signalling blocking conditions. Tracks show counts normalised to RPKM of two merged biological replicates.

**Supplementary Fig. 1.**
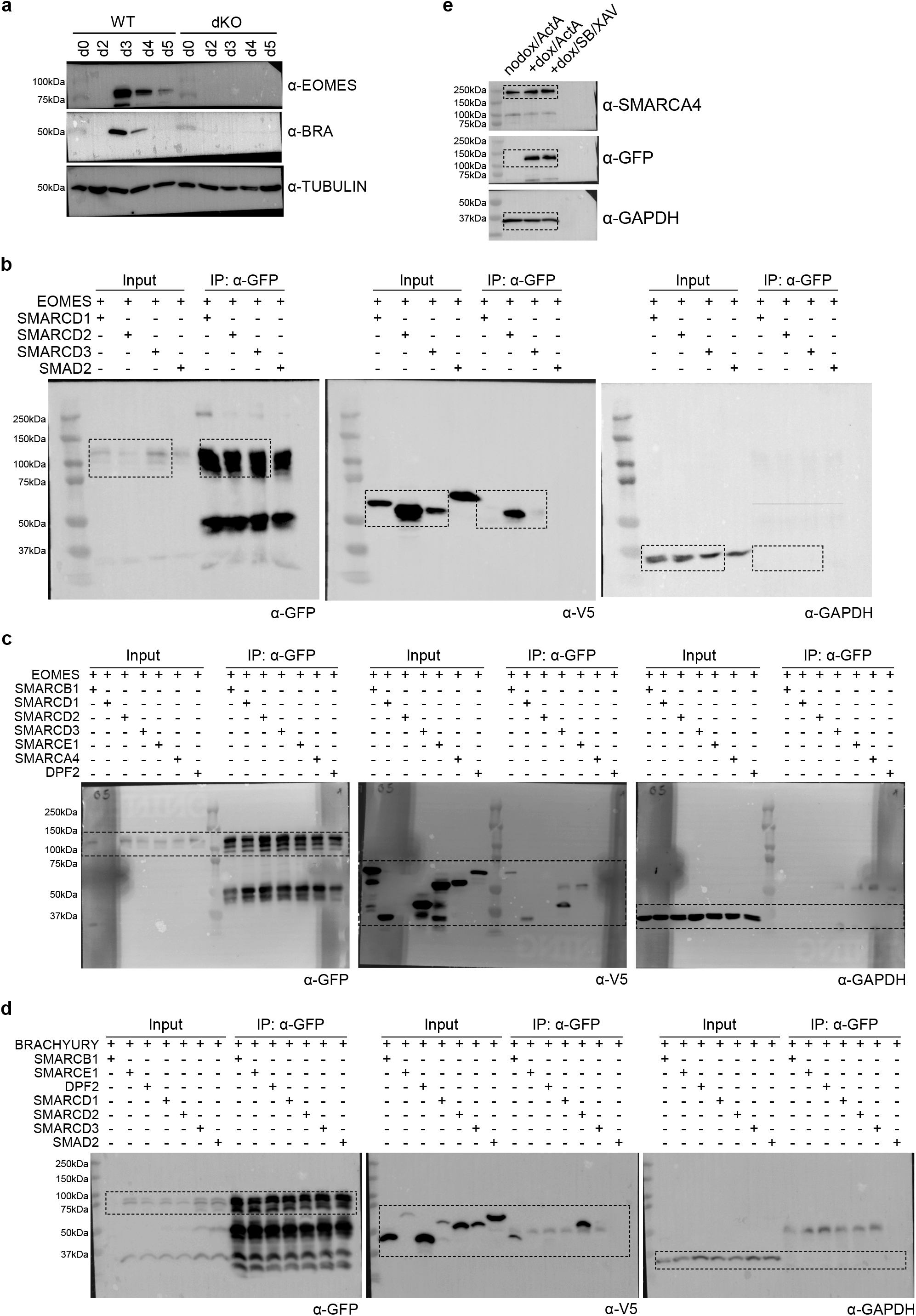
Unprocessed immunoblots. **a**, Unprocessed image of Western blot relates to Fig. 3b; **b**, relates to Fig. 4e; **c**, relates to Extended Data Fig. 5a; **d**, relates to Extended Data Fig. 5b; **e**, relates to Extended Data Fig. 6b.

